# Regulation of Iron Homeostasis by SPAK-dependent Modulation of FBXL5 Stability

**DOI:** 10.1101/133777

**Authors:** David N. Powers, Ajay A. Vashisht, Monika Shenouda, Calvin Yao, James A. Wohlschlegel

## Abstract

Intracellular iron homeostasis is regulated by a proteolytic switch whereby the E3 ubiquitin ligase FBXL5 targets iron regulatory proteins (IRPs) for ubiquitin-dependent degradation in iron-replete conditions while it is itself degraded during iron deficiency allowing IRPs to accumulate and regulate their downstream RNA targets. The cellular pathways that control FBXL5 degradation in low iron conditions are not well understood. Here, we report the identification of the STE20/SPS1-related proline-alanine-rich protein kinase (SPAK) as a novel regulator of FBXL5 stability. We find that SPAK, a kinase previously implicated in osmotic stress regulation, regulates intracellular iron homeostasis through its physical association with FBXL5. This role is dependent on its kinase activity as overexpression of constitutively active SPAK increases FBXL5 poly-ubiquitination and degradation. Through this work, we have discovered a novel role for SPAK that extends beyond its well-established function in salt homeostasis and raises the possibility for signaling crosstalk between iron homeostatic and osmotic regulatory pathways.

## INTRODUCTION

Iron is an essential metal cofactor due to its ability to accept and donate electrons - it is this reactivity that results in its utilization in numerous biochemical processes such as oxygen delivery and storage, DNA repair and replication, lipid metabolism and chromatin modification (1). Abnormalities in iron homeostasis result in numerous human diseases such as anemia, hereditary hemochromatosis and aceruloplasminemia (2). Due to the importance of proper iron maintenance at both the cellular and systemic level, the mechanisms of iron acquisition, distribution and regulation are major avenues of research.

Despite the critical role of iron in the cell, many aspects of its intracellular regulation remain unknown. We and others previously discovered an E3 ubiquitin ligase FBXL5 that regulates the ubiquitin-mediated degradation of the mRNA-binding proteins IRP1 and IRP2 in an iron-dependent manner that is controlled by iron binding directly to a hemerythrin-like domain present in its N-terminus (3-5). FBXL5 is stabilized by iron binding and assembles into an E3 ubiquitin ligase complex that targets IRP1 and IRP2 for degradation; without iron, FBXL5 is itself degraded. The mechanism for FBXL5 degradation is not well characterized.

In this paper, we identify a novel interaction between FBXL5 and SPAK, a STE20/SPS1-related proline-alanine-rich protein kinase involved in osmotic regulation. We report that SPAK influences FBXL5 stability leading to downstream alterations in iron homeostatic pathways. This effect is dependent on SPAK activity with constitutively active and kinase dead SPAK mutants having distinct effects on FBXL5 function. Our results open up new connections between cellular stress response pathways that have, until now, been largely separated.

## EXPERIMENTAL PROCEDURES

*Plasmids -* The full length FBXL5 cDNA (NIH_MGC_97) and full length mouse SPAK cDNA (Clone ID: 5698326) were obtained from Open Biosystems. The cDNA was amplified using the Phusion DNA Polymerase (New England Biolabs) and introduced into the pCR8/GW/TOPO vector (Invitrogen) or the Gateway pDONR221 vector (Life Technologies). The Quikchange system (Stratagene) was used to generate the FBXL5-ΔFbox mutant lacking amino acids 216-240 using pCR8-FBXL5 as a template and was used to create the substitution mutations in FBXL5 and SPAK (K104R, T243E/S383D) using a pDEST-SPAK-WT vector as a template. FBXL5 and FBXL5-ΔFbox were subcloned into pcDNA3-6xMyc, pcDNA5-FRT/TO-3xHA-3xFLAG and pMT/BioEase^™^-DEST plasmids using pCR8-FBXL5 and DEST plasmids via the Gateway cloning system (Life Technologies). FBXL5 and SPAK fragments were generated by PCR using primers containing flanking AttB1 and AttB2 sites and cloned into pDONR221. These fragments were then subcloned into pcDNA5-FRT/TO-3xHA-3xFLAG for expression in HEK293 or HeLa. Plasmids expressing HA-Ub were previously described (6,7).

*Antibodies -* Antibodies used for immunoblotting were as follows: IRP2-7H6 (Santa Cruz Biotechnology), HERC2 (Bethyl Laboratories, Inc), FBXL5 (neoclone), SPAK A302-466A (Bethyl Laboratories, Inc), FLAG-M2 (Sigma), c-Myc (Santa Cruz Biotechnology), ferritin (Sigma), β-tubulin (Sigma) and α-tubulin-HRP (Proteintech Group). Streptavidin-HRP (Life Technologies) was used to detect BioEase-tagged proteins. Horseradish peroxidase conjugated secondary antibodies were obtained from Jackson Immunoresearch Laboratories. Immunoprecipitation reactions were performed using affinity matrices for anti-FLAG M2, anti-c-Myc, and anti-HA available from Sigma.

*Cell Lines -* The HEK293 and HeLa cell lines were obtained from the American Type Culture Collection (ATCC) while Flp-In^TM^ T-REx^TM^-293 was obtained from Invitrogen. Flp-In^TM^ T-REx^TM^-293 cells stably expressing 3xHA-3xFLAG-FBXL5, 3xHA-3xFLAG-FBXL5-ΔFbox, 3xHA-3xFLAG-FBXL5-FB2, 3xHA-3xFLAG-FBXL5-FB2-ΔFbox, 3xHA-3xFLAG-SPAK, 3xHA-3xFLAG-SPAK-K104R, 3xHA-3xFLAG-SPAK-T243E/S383D were generated using the Flp-In system (Invitrogen) according to the manufacturer’s directions.

*Cell Culture, Plasmid Transfections, and Treatments -* Cell culture reagents were obtained from Invitrogen. All cell lines were cultured in complete Dulbecco's Modified Eagles medium (DMEM) containing 10% heat inactivated fetal bovine serum (FBS), 100 units/mL penicillin and streptomycin, and 2 mM glutamine at 37**°**C in ambient air with 5% CO2. Transient transfections were performed using either BioT (Bioland, Long Beach, CA) or Lipofectamine 2000 according to the manufacturer’s protocol. siRNA transfections were performed according to the manufacturer’s protocol using Lipofectamine RNAiMAX (Life Technologies) and siGENOME SMARTpool reagents for HERC2 (Dharmacon #M-007180-02), SPAK (Dharmacon #M-004875-02-0005) or a non-targeting siGENOME control siRNA (Dharmacon #D-001210-03-05). Expression of 3xHA-3xFLAG-tagged protein in a stable cell line was induced by treating cells with doxycycline for 24 hours at a final concentration of 500 ng/mL. Cells were treated with 100 μg/ml ferric ammonium citrate (FAC) (Thermo Fisher), 100 μM desferrioxamine mesylate (DFO) (Sigma), and/or or 25 μM MG132 (Z-Leu-Leu-Leu-CHO) (BIOMOL) for the times indicated.

*Affinity purification of FBXL5-ΔFbox Protein Complexes -* Twenty-five 15-cm tissue culture plates of Flp-In^TM^ TREx^TM^-293 cells stably expressing His6-3xFLAG-FBXL5-ΔFbox were grown, harvested, and lysed in IP buffer (100 mM Tris-HCl pH 8.0, 150 mM NaCl, 5 mM EDTA, 5% glycerol, 0.1% NP-40, 1 mM DTT, 0.5 mMPMSF, 1 μM pepstatin, 1 μM leupeptin and 2 μg/mL aprotinin). 200-300 mg of clarified protein lysate was then incubated at 4°C with 100 μL of equilibrated anti-FLAG M2 agarose (Sigma) for 2 hours. Beads were then washed four times using 1 mL of IP Buffer per wash before eluting with 500 μL of FLAG Elution Buffer (IP buffer lacking NP-40 and supplemented with 250 μg/mL of 3xFLAG peptide (Sigma)). Elutions were precipitated by the addition of trichloroacetic acid (TCA) to a final concentration of 10% followed by incubation on ice for 30 minutes and centrifugation at 16,000g for 10 minutes to collect the precipitate

*Proteomic Characterization of FBXL5 purifications -* TCA precipitates from affinity-purified His6-3xFLAG-FBXL5-ΔFbox cells were digested and prepared for proteomic analysis as described (8). The digested samples were analyzed by MudPIT (9,10). A 5-step multidimensional chromatographic separation was performed online and fractionated peptides were eluted directly into a LTQ-Orbitrap XL mass spectrometer (Thermo Fisher) in which tandem mass spectra were collected. Peptide mass spectra were analyzed using the SEQUEST and DTASelect algorithms (11,12). A decoy database approach was used to estimate peptide and protein level false positive rates which were less than 5% per analysis (13). Proteins were considered candidate FBXL5 interacting proteins if they were identified in the relevant affinity purification but not in MudPIT analyses of other control purifications. A detailed description of the multidimensional peptide fractionation protocol, mass spectrometer settings, and bioinformatic workflow is described elsewhere (14).

*Immunoprecipitation and Immunoblotting* **-** Cell lysates were prepared using IP buffer with Simple Stop^TM^ 2 Phosphatase Inhibitor Cocktail (Gold Biotechnology). Immunoprecipitations were performed using the appropriate affinity matrix equilibrated with IP Buffer and incubated with equal amounts of cell lysates at 4°C for 2 hours. Beads were washed three times with wash buffer (IP buffer without protease/phosphatase inhibitors) and resuspended in 2x SDS loading buffer. For immunoblotting, whole cell lysates and immunoprecipitates were boiled in SDS-loading buffer, separated using SDS-PAGE, transferred to Immobilon-P or Immobilon-FL PVDF membranes (Millipore), and probed with the appropriate primary and secondary antibodies. Proteins were visualized using Pierce ECL western blotting substrate (Thermo Fisher) or the LI-COR Odyssey Imager

*Ubiquitination assay -* HEK293 cells were transfected with plasmids expressing HA-Ubiquitin, a 3xFLAG FBXL5 construct (3xFLAG-FBXL5, 3xFLAG-FBXL5-ΔFbox), and combinations of 6xMyc-SPAK-K104R, 6xMyc-SPAK-T243E/S383D and 6xMyc-SPAK-T243E/S383DΔCCT. Twenty-four hours after transfection, the medium was changed and cells were treated with 100μg/ml FAC and 25 μM MG132 for 4 hours. Cells were harvested and lysed under denaturing conditions as described previously (15). Ubiquitin conjugates were purified using anti-HA beads and the presence of FBXL5 in the purified ubiquitin conjugates was detected by immunoblotting with FLAG M2 antibody.

*In Vitro Binding Competition Assay -* HEK293 cells in a 6-well plate were transfected with either BioEase-FBXL5, 3HA-3FLAG-tagged SPAK or SKP1. The cells were harvested separately and lysed in IP buffer. BioEase-FBXL5 was purified using High Capacity Streptavidin Agarose Resin (Thermo Scientific), while 3HA-3FLAG-tagged SPAK and SKP1 were bound to EZview Red ANTI-FLAG M2 Affinity Gel (Sigma) 1-hour binding at 4°C on rotisserie rotator. Beads were washed 3 times with wash buffer. SPAK and SKP1 were eluted off of the ANTI-FLAG Gel using two 20 minute washes of 250μg/mL FLAG peptide at 4°C on rotisserie rotator. Eluted SPAK and SKP1 were incubated with FBXL5-bound Agarose Resin for 1 hour at 4°C on rotisserie rotator, and then washed 3 times with wash buffer. The second incubation step with either SKP1 or SPAK with FBXL5-bound Agarose Resin took place for 1 hour at 4^o^C on rotisserie rotator, and then washed 3 times with wash buffer prior to addition of SDS loading dye and boiling at >100°C for 3 minutes.

## RESULTS

To identify novel cellular factors that regulate FBXL5 stability, proteomic mass spectrometry was used to identify proteins that co-purify with FBXL5 using an affinity-purification mass spectrometry (AP-MS) approach. Briefly, FBXL5-associated protein complexes were immuno-purified from an HEK293 stable cell line expressing HA-FLAG-FBXL5 from a doxycycline-inducible promoter. These FBXL5 immunoprecipitates (IP) were digested with trypsin and analyzed by Multidimensional Protein Identification Technology (MudPIT) using an LTQ-Orbitrap XL tandem mass spectrometer (3,5). Purifications were also done using a dominant negative variant of FBXL5 (FBXL5-ΔFbox) which has a portion of its F-box region deleted (amino acids 216 – 240) in order to increase its ability to trap potential substrate proteins (16). In addition to previously reported IRPs (3), this proteomic analysis identified three additional partners of FBXL5 that function in the cellular osmotic stress response: WNK1, SPAK and OSR1 (Figure 1A). WNK1, SPAK and OSR1 make up a kinase cascade pathway that regulates osmotic stress. This pathway consists of upstream WNK (With No-[K] Lysine) kinases (WNK1, WNK2, WNK3 and WNK4) which regulate SPAK activation and the phosphorylation of downstream targets such as the membrane-expressed Na^+^-K^+^-Cl^-^ Cotransporter 1 (NKCC1) (17). SPAK is phosphorylated by activated WNK1 in conditions of osmotic stress – for example, a hyperosmotic treatment such as sorbitol or hypotonic low-Cl^-^ conditions (18,19). Activated SPAK phosphorylates and activates NKCC1 to transport salt into the cell (20). All of SPAK’s known interacting partners including its upstream kinases and downstream substrates associate with SPAK through its conserved C-Terminal (CCT) domain (21). SPAK binding partners such as WNK1 and NKCC1 feature RFxV motifs that are required to bind the CCT domain.

**FIGURE 1.**
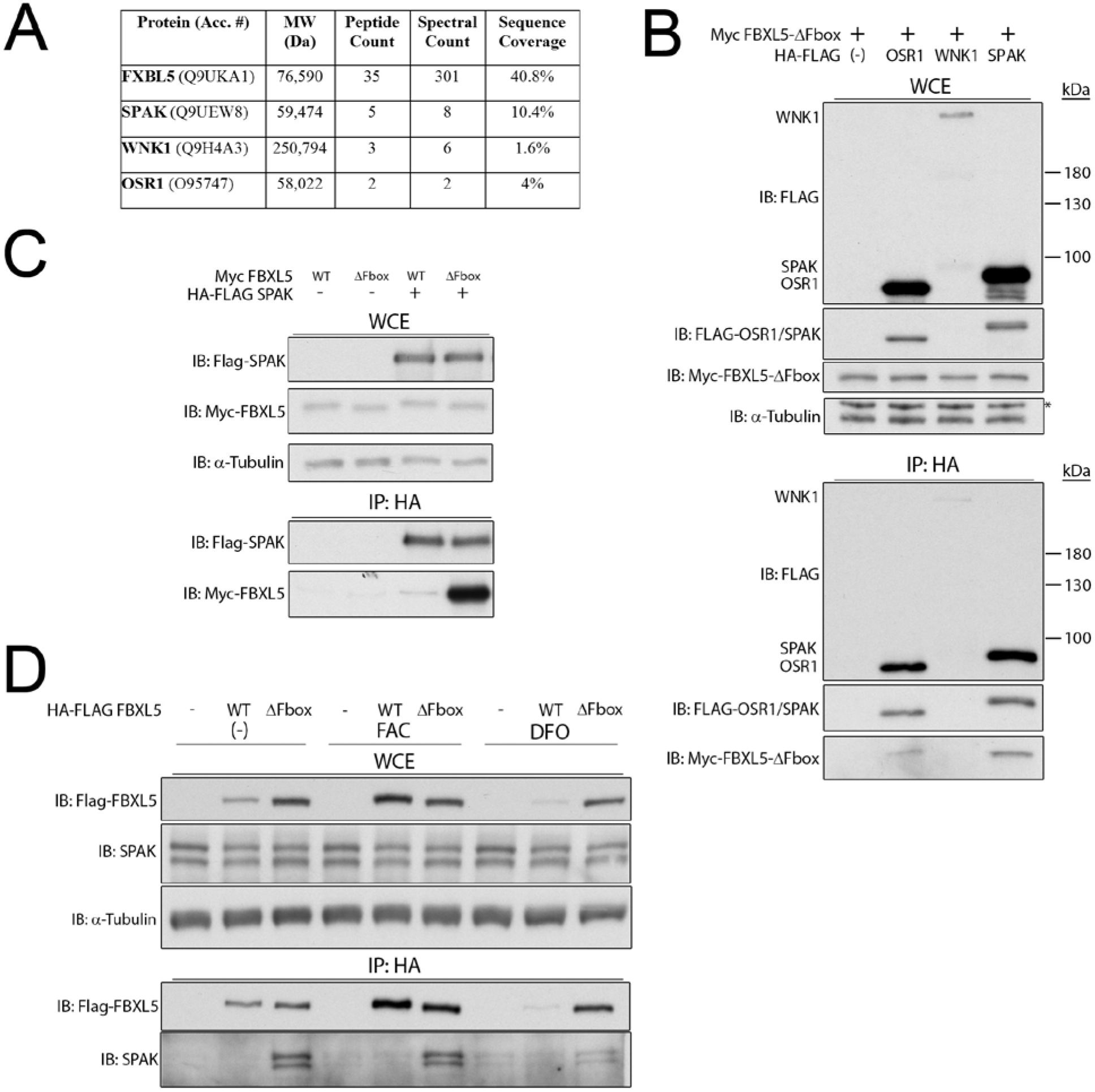
SPAK interacts with FBXL5 and FBXL5-ΔFbox. (**A**) Binding partners identified in the proteomic analysis of FBXL5-ΔFbox. (**B**) HEK293 cells were transiently transfected with 6Myc-FBXL5-ΔFbox and 3HA-3FLAG-tagged OSR1, WNK1 or SPAK. HA-tagged proteins and their interacting partners were immunopurified using anti-HA beads and the whole cell extracts (WCE) and immunoprecipitations (IP) were blotted with primary antibodies against FLAG, c-Myc and α-tubulin. (**C**) HEK293 cells were transiently transfected with combinations of 6Myc-tagged FBXL5-WT or FBXL5-ΔFbox together with 3HA-3FLAG-SPAK. SPAK was affinity purified using anti-HA beads and the WCEs and IPs were blotted with primary antibodies against FLAG, c-Myc and α-tubulin. (**D**) HEK293 cells stably expressing either 3HA-3FLAG-tagged FBXL5 wild type (WT) or FBXL5-ΔFbox were affinity-purified using antibodies against HA from cells treated with 8 hours of FAC or DFO as indicated. WCEs and IPs were blotted against with antibodies for FLAG, SPAK and α-tubulin.

Oxidative Stress-Responsive 1 protein (OSR1) is a close structural homologue of SPAK. These two kinases share very high sequence homology: 96% in the N-terminal catalytic region and 67% in the C-terminal regulatory domain (22). The region that differs the most between the two proteins is a unique proline-alanine rich 48 amino acid long stretch in the N-terminus of SPAK from which the protein derives its name. The function of this proline-alanine rich region is still unknown, as SPAK can function normally as a kinase without it (23). In spite of the high degree of similarity between SPAK and OSR1, mouse knockout studies revealed that the disruption of SPAK results in viable animals with neurobehavioral differences from wild type adults, while OSR1 knockout causes embryonic lethality and generates no living pups (22,24). This indicates that while SPAK and OSR1 appear to be functionally similar within the osmotic stress regulatory pathway, important functional differences remain to be elucidated.

To validate the AP-MS results, we characterized the SPAK-FBXL5 interaction using biochemical methods. We transiently co-transfected Myc-tagged FBXL5-ΔFbox together with HA-FLAG-tagged WNK1, OSR1 and SPAK in HEK293 cells and determined the ability of the proteins to interact using co-immunoprecipitation (co-IP) followed by Western blotting (Figure 1B). Of the three kinases, FBXL5-ΔFbox interacted most strongly with SPAK, so we focused our efforts on characterizing the interaction between these two proteins. Next, we transiently expressed Myc-tagged versions of FBXL5 and FBXL5-ΔFbox with HA-FLAG-tagged SPAK and used co-IP assays to examine the putative interaction between SPAK and wild-type FBXL5 (FBXL5-WT). As seen in Fig. 1C, we observed that the SPAK co-purifies with both FBXL5-WT as well as FBXL5-ΔFbox with the SPAK–FBXL5-ΔFbox interaction appearing to be the significantly stronger of the two. We also demonstrated that FBXL5-ΔFbox associates with endogeneous SPAK in an iron-regulated manner (Figure 1D). Specifically, SPAK is detected in FBXL5-ΔFbox immunoprecipitates from cells treated with ferric ammonium citrate (FAC) which results in high-iron conditions but the interaction is much weaker in cells treated with the iron chelator desferrioxamine mesylate (DFO). It is worth noting that FBXL5-ΔFbox is not degraded during iron deficiency and yet still loses its interaction with SPAK in low iron, suggesting that this iron-regulated interaction works directly on the level of the FBXL5–SPAK interaction and not indirectly through modulation of FBXL5 levels.

After discovering that the interaction between FBXL5 and SPAK is iron-dependent, we hypothesized that SPAK might influence cellular iron homeostasis. We find that depletion of SPAK by siRNA leads to the accumulation of endogenous FBXL5 as well as changes in the stability of downstream factors, iron-regulatory proteins IRP2 and ferritin (Figure 2A). As a positive control we have included results for the siRNA knockdown of HERC2, an E3 ubiquitin ligase known to target FBXL5, which also leads to the accumulation of IRP2 and ferritin (25).

**FIGURE 2.**
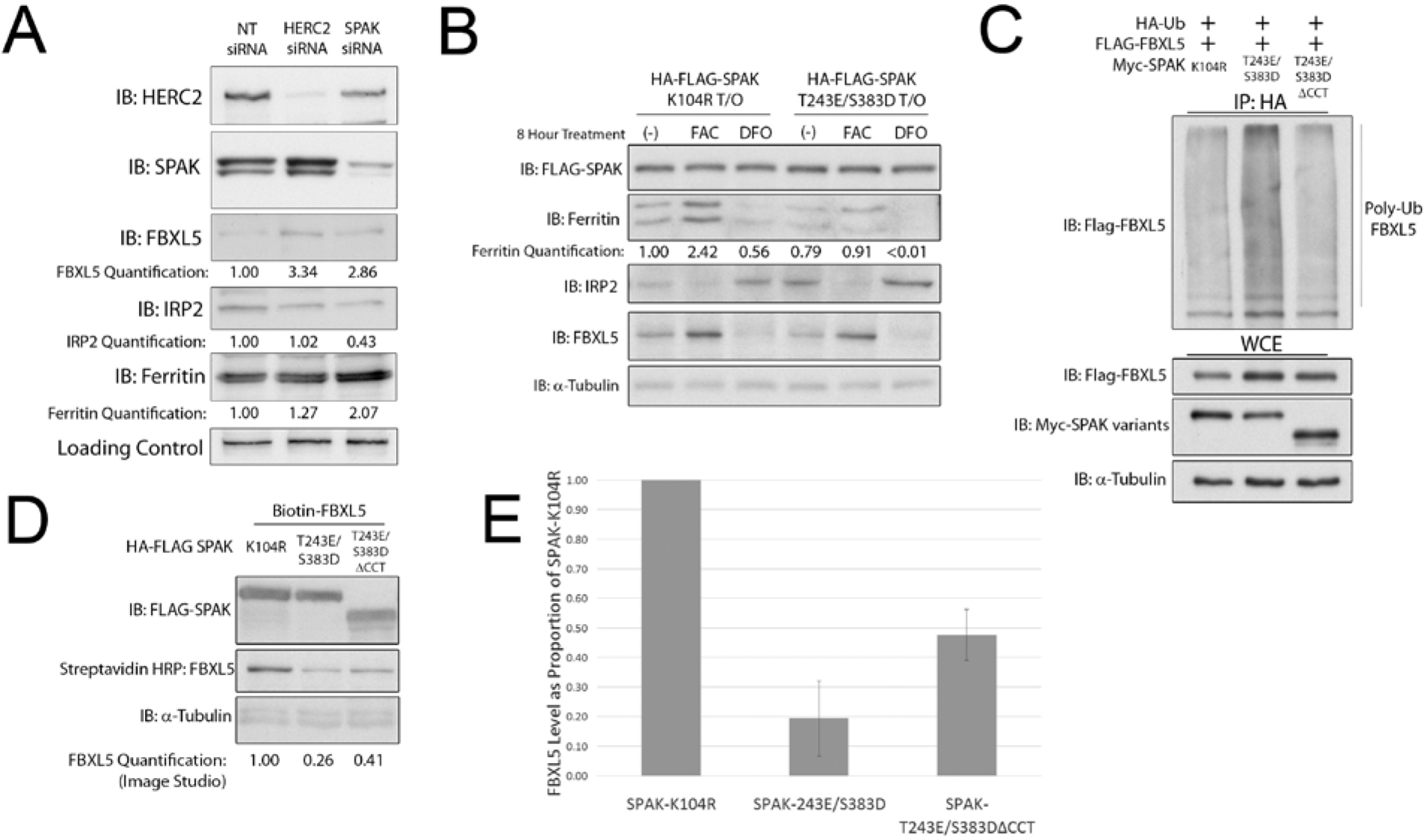
SPAK depletion or expression of SPAK mutants affects the stability of iron-regulatory proteins. (**A**) Transfection of siRNA duplexes directed against HERC2 or SPAK were used to silence their expression in HEK293 cells. WCEs were probed with primary antibodies against HERC2, SPAK, FBXL5, IRP2 and ferritin. Relative protein signals were quantified using the Image Studio software. (**B**) HEK293 stable cell lines expressing 3HA-3FLAG tagged SPAK-K104R or SPAK-T243E/S383D were treated for 8 hours with FAC or DFO prior to lysis. WCEs were blotted with antibodies against FLAG, ferritin, IRP2, FBXL5 and α-tubulin. (**C**) HEK293 cells were cotransfected with HA-Ub and FLAG-FBXL5 together with either Myc-SPAK-K104R, Myc-SPAK-T243E/S383D or Myc-SPAK-T243E/S383D-ΔCCT and treated with MG132 for 4 hours. Ubiquitin conjugates were immunoprecipitated using anti-HA beads. HA-immunoprecipitates (IP: HA) and WCEs were immunoblotted with antibodies to FLAG, c-Myc, and α-tubulin. (**D**) HEK293 cells were transiently transfected with BioEase-FBXL5 and 3HA-3FLAG-tagged variants of SPAK: SPAK-K104R, SPAK-T243E/S383D or SPAK-T243E/S383D-ΔCCT. WCEs were blotted with streptavidin-HRP and primary antibodies to FLAG and α-tubulin. FBXL5 levels were quantitated using the Image Studio software. (**E**) Graph representing results from 4 biological replicates of BioEase-FBXL5 destabilization by SPAK-T243E/S383D and SPAK-T243E/S383DΔCCT. Confidence intervals are as calculated for Student’s t-test with α=0.05 and *n*=4.

Based on these observations that SPAK can influence intracellular iron homeostasis, we next examined whether SPAK kinase activity was important for this function. We generated doxycycline inducible stable cell lines that either express kinase-dead (SPAK-K104R) or constitutively-active (SPAK-T243E/S383D) mutant forms of SPAK. We observed that overexpression of SPAK-K104R was able to stabilize endogenous levels of FBXL5 leading to downregulation of IRP2 and depression of IRP2-mediated ferritin translation (Figure 2B). In particular, steady state ferritin levels are considerably higher in SPAK-K104R overexpressing cells as compared to SPAK-T243E/S383D overexpressing conditions (Figure 2B, compare lanes 1 & 2 with lanes 4 & 5).

Given that SPAK knock-down and overexpression of SPAK-K104R increase stabilization of FBXL5, we reasoned that SPAK may modulate ubiquitin-dependent degradation of FBXL5 in a kinase activity-dependent manner. To test this possibility, we examined whether overexpression of either SPAK-T243E/S383D or SPAK-K104R could stimulate FBXL5 ubiquitination in a cell-based ubiquitination assay that measures the accumulation of FBXL5 in affinity-purified ubiquitin conjugates after treatment with the proteasome inhibitor MG132. In Figure 2C, we find that overexpression of SPAK-T243E/S383D was able to stimulate FBXL5 poly-ubiquitination while SPAK-K104R could not. Moreover, the ability of SPAK-T243E/S383D to stimulate FBXL5 ubiquitination was dependent on the presence its CCT domain, a conserved domain present in its C-terminus that has previously been shown to mediate SPAK interactions with upstream and downstream signaling components. Consistent with these cell-based ubiquitination results, we also find that overexpression of active SPAK but not kinase dead SPAK leads to a reduction in steady-state FBXL5 levels and that this effect was also at least partially dependent on its CCT domain (Figure 2D, E). Together, these results indicate that constitutively active SPAK-T243E/S383D is able to stimulate the poly-ubiquitination and subsequent degradation of FBXL5 in a manner dependent on its CCT domain. We hypothesize that the CCT domain of SPAK is responsible for recruiting an uncharacterized protein that can result in conditions that promote FBXL5 poly-ubiquitination and degradation.

To further characterize the FBXL5–SPAK interaction, we conducted domain mapping experiments to identify the regions in both FBXL5 and SPAK that mediate this interaction. We reasoned that this data would help us establish the biological role of the interaction. To map the region of FBXL5 that mediates its interaction with SPAK, we generated a series of deletion mutants of FBXL5 and tested their ability to associate with SPAK by co-IP in order to identify the minimal binding region required for the interaction (Figure 3A). We find that a C-terminal 532 amino acid fragment of FBXL5-ΔFbox (FBXL5-ΔFbox-CT532) lacking both the hemerythin-like domain and most of the F-box domain retains high SPAK binding. However, a smaller C-terminal fragment containing only the substrate-binding leucine rich repeats (FBXL5-CT443) loses the ability to bind to SPAK. This suggests that SPAK associates with the region between the hemerythrin-like domain and the F-box domain of FBXL5 that spans amino acids 159 to 215. It also indicates that SPAK is likely not a substrate of FBXL5 since the SPAK interaction does not map to the substrate binding motif of FBXL5. To further define the important binding elements in FBXL5 region amino acids 159 to 215, we created additional FBXL5-ΔFbox N-terminal truncation mutants within this region of the protein and tested their ability to associate with SPAK-WT (Figure 3B). While the C-terminal 452 amino acids of FBXL5 (FBXL5-CT452) were unable to strongly bind SPAK, the addition of amino acids 202 to 215 to the N-terminal end of this fragment (FBXL5-ΔFbox-CT490) restored strong SPAK binding. This indicates that the SPAK binding motif is likely found within the FBXL5 sequence: STGITHLPPEVMLS.

**FIGURE 3.**
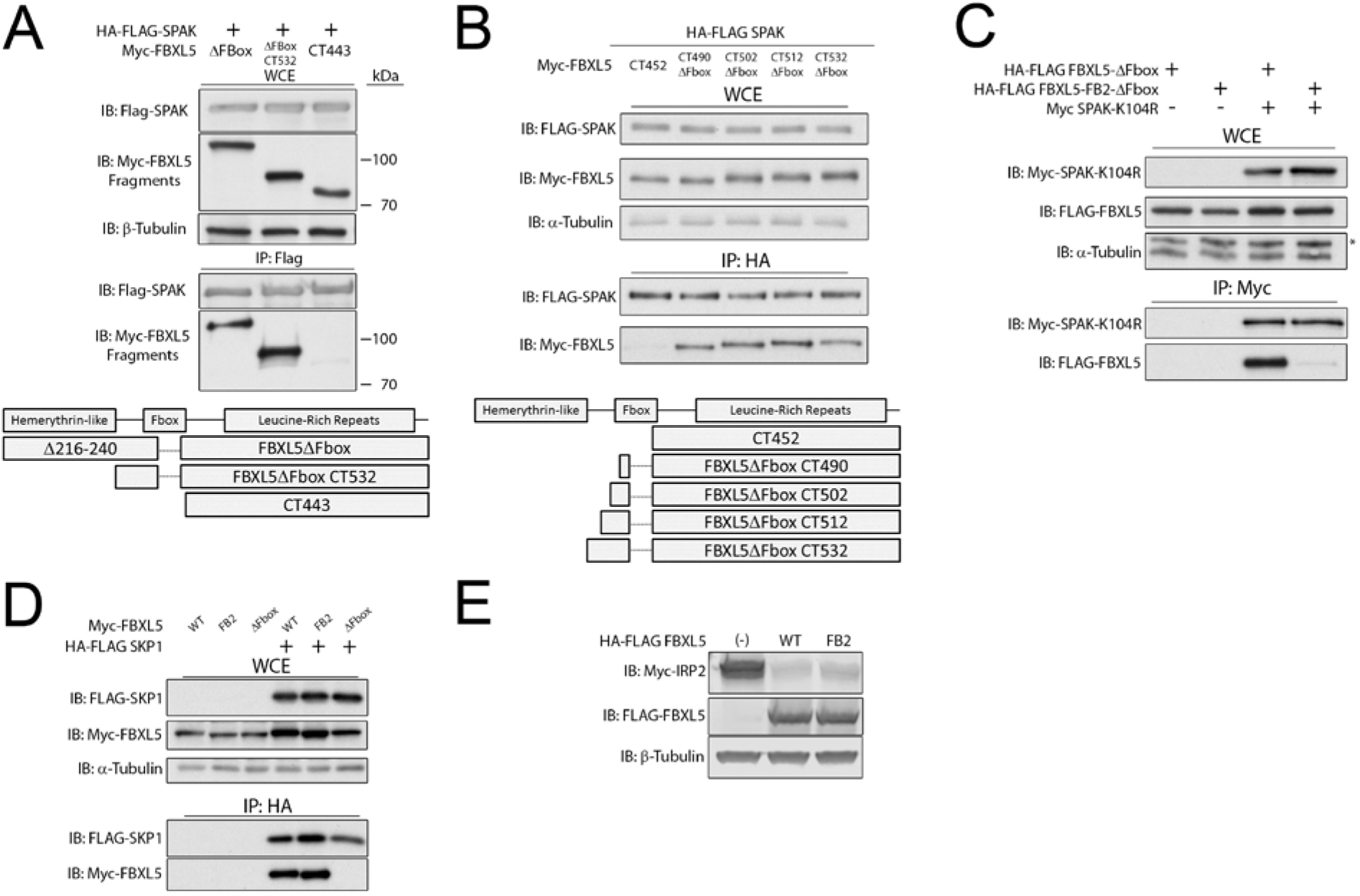
Characterizing the SPAK–FBXL5 interaction. (**A**) 3HA-3FLAG SPAK was transiently transfected into HEK293 cells with 6Myc-tagged fragments of FBXL5. FLAG affinity purification was used against FLAG-SPAK. WCE and IP were blotted with FLAG, Myc and α-Tubulin primary antibodies. (**B**) 3HA-3FLAG SPAK was transiently transfected into HEK293 cells with 6Myc-tagged fragments of FBXL5-ΔFbox. HA affinity purification was used against HA-SPAK. WCE and IP were blotted with FLAG, c-Myc and α-Tubulin primary antibodies. (**C**) 3HA-3FLAG-tagged FBXL5-ΔFbox and FBXL5-FB2-ΔFbox were co-transfected with 6Myc-tagged SPAK-K104R. SPAK-K104R and its complexes were immunoprecipitated with anti-Myc beads, and the WCE and IP were blotted with FLAG, c-Myc and α-Tubulin. (**D**) 3H3F-tagged SKP1 was co-transfected with 6Myc-tagged variants of FBXL5, as indicated. SKP1 complexes were immunoprecipitated using HA beads. WCE and IP were Western blotted with antibodies against FLAG, c-Myc and α-Tubulin. (**E**) HEK293 cells were co-transfected with Myc-tagged IRP2 and either 3HA-3FLAG-tagged FBXL5 or FBXL5-FB2. WCEs were collected and Western blotted with primary antibodies against c-Myc, FLAG and β-tubulin.

As the STGITHLPPEVMLS sequence is adjacent to the F-box domain, we were concerned that mutations in this region might be affecting SPAK binding indirectly by influencing SKP1 binding. To address this potential problem, we substituted this 14 amino acid stretch in FBXL5 with the analogous region from the F-box protein FBXL2 (GLINKKLPKELLLR) in order to create a FBXL5 mutant that retains SKP1 binding but was still defective in SPAK binding. This mutant features amino acid substitutions at 10 amino acids across the 14 amino acid stretch (**STGITH***LP***P***E***VM***L***S**, substituted amino acids are **bolded**), which we denoted as FBXL5-FB2 (FBXL5 Fbox 2). When the ΔFbox variant of FBXL5-FB2 (denoted as FBXL5-FB2-ΔFbox) was co-expressed with SPAK-K104R, we observed that FBXL5-FB2-ΔFbox was unable to interact via co-IP (Figure 3C). We used SPAK-K104R instead of SPAK-WT to create conditions with the strongest interaction and show that binding is essentially eliminated when incorporating the FB2 mutation. As predicted, FBXL5-FB2 retained SKP1 binding ability (Figure 3D). Together, these results identify a region in FBXL5 that is N-terminal to the F-box domain and is required for SPAK binding

To validate that FBXL5-FB2 retains its function as an E3 ubiquitin ligase, we measured its ability to stimulate IRP2 degradation in a transient transfection assay (Figure 3E). We observed that overexpression of FBXL5-WT and FBXL5-FB2 both drive degradation of co-transfected IRP2, indicating that FBXL5-FB2 is functional and can promote the ubiquitination and degradation of IRP2.

We next tested whether loss of SPAK binding by FBXL5 would affect either its stability or the stability of its downstream substrates. In cell lines stably expressing either 3HA-3FLAG-FBXL5-WT or FBXL5-FB2, we observed that loss of SPAK binding led to a reduction in FBXL5 abundance and a corresponding increase and decrease in levels of IRP2 and ferritin, respectively. (Figure 4A). These results indicate the importance of the SPAK interaction for maintaining the stability of FBXL5 and its regulation of downstream signaling components. We observed that FBXL5-FB2 was present at lower protein levels relative to FBXL5-WT in these cell lines. To determine if this is due to the reduced stability of FBXL5-FB2, we determined the half-life of FBXL5-FB2 using cycloheximide (CHX) treatment to block protein synthesis. We discovered that the half-life of FBXL5-FB2 is significantly lower than FBXL5-WT (1.8 hours versus 2.7 hours), indicating that SPAK binding regulates FBXL5 degradation (Figure 4B). Additionally, this degradation is proteasome-dependent, as treatment with the proteasome inhibitor MG132 blocks the increased degradation seen in the FBXL5-FB2 relative to FBXL5-WT (Figure 4C). Together, these results support a model in which SPAK regulates the proteasome-dependent degradation of FBXL5.

**FIGURE 4.**
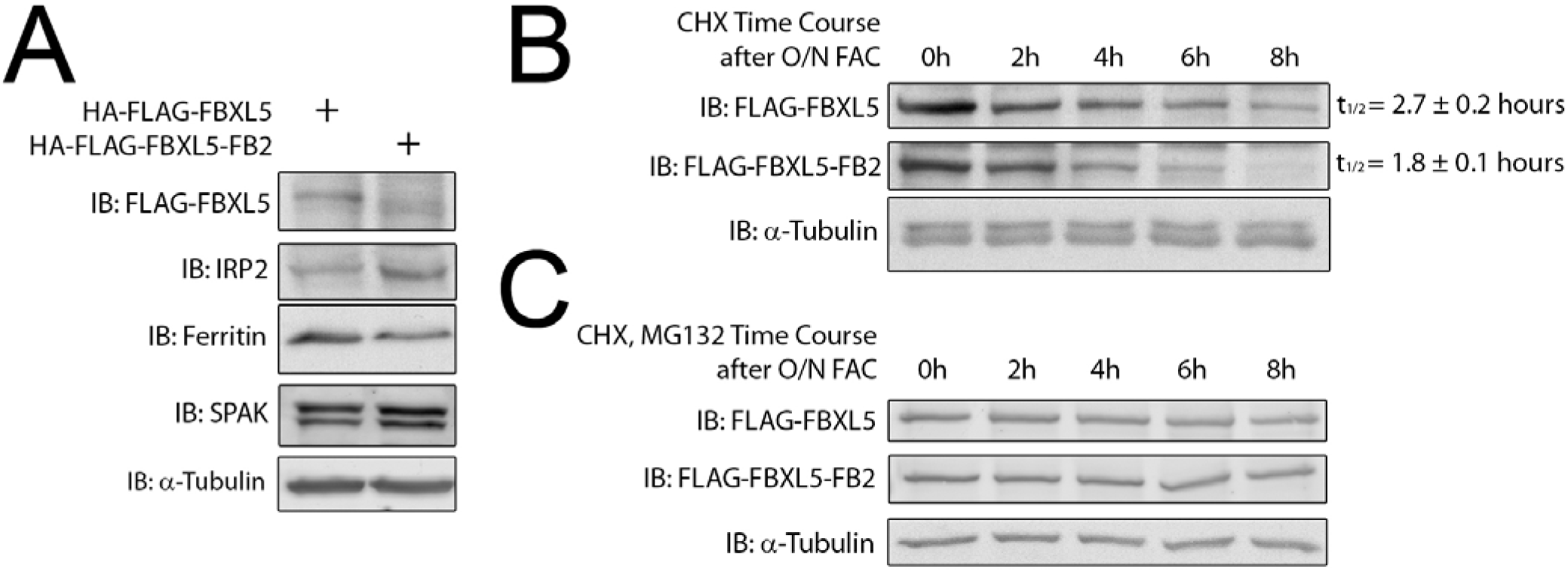
Loss of SPAK binding decreases the stability of FBXL5 and affects the stability of downstream iron-regulatory proteins. (**A**) Stable cell lines expressing 3HA-3FLAG-tagged FBXL5 or FBXL5-FB2 were lysed and the WCEs Western blotted with primary antibodies against FLAG, IRP2, ferritin, SPAK and α-tubulin. (**B**) HEK293 cells stably expressing 3H3F-tagged FBXL5 or FBXL5-FB2 were treated overnight with doxycycline and FAC. The media was changed 18 hours later and cycloheximide added at the designated time-points. WCEs were blotted with primary antibodies against FLAG and α-tubulin. The half-life calculation was based on Image Studio quantification. (**C**) HEK293 cells stably expressing 3HA-3FLAG-tagged FBXL5 or FBXL5-FB2 were treated overnight with doxycycline and FAC. The media was changed 18 hours later and cycloheximide with MG132 was added at the designated times. WCEs were blotted with primary antibodies against FLAG and α-tubulin.

To further explore the mechanism by which SPAK is regulating FBXL5 stability, we examined the FBXL5–SKP1 interaction. Since SPAK and SKP1 appear to bind to overlapping regions of FBXL5, we hypothesized that they might compete for FBXL5 binding and thereby regulate its activity and/or stability. Consistent with this idea, the co-expression of FBXL5 and SPAK-K104R with increasing concentrations of SKP1 led to stabilization of FBXL5 in a SKP1-dependent manner (Figure 5A). In addition, we used an *in vitro* binding assay to directly assess whether SPAK and SKP1 compete for FBXL5 binding. Biotinylated FBXL5 was bound to streptavidin beads and then incubated with immunopurified HA-FLAG-tagged SPAK or SKP1. After the initial incubation, unbound protein was washed away and the beads were then re-incubated with the other factor (i.e. SPAK then SKP1 or vice-versa). In both cases, we observed that SPAK and SKP1 were able to displace each other from FBXL5 (Figure 5B) and that this displacement occurred in a concentration-dependent manner (Figure 5C). We conclude that SPAK and SKP1 share an overlapping binding site on FBXL5 that is mutually exclusive, and that binding to either SPAK or SKP1 differentially regulates FBXL5 function and stability.

**FIGURE 5.**
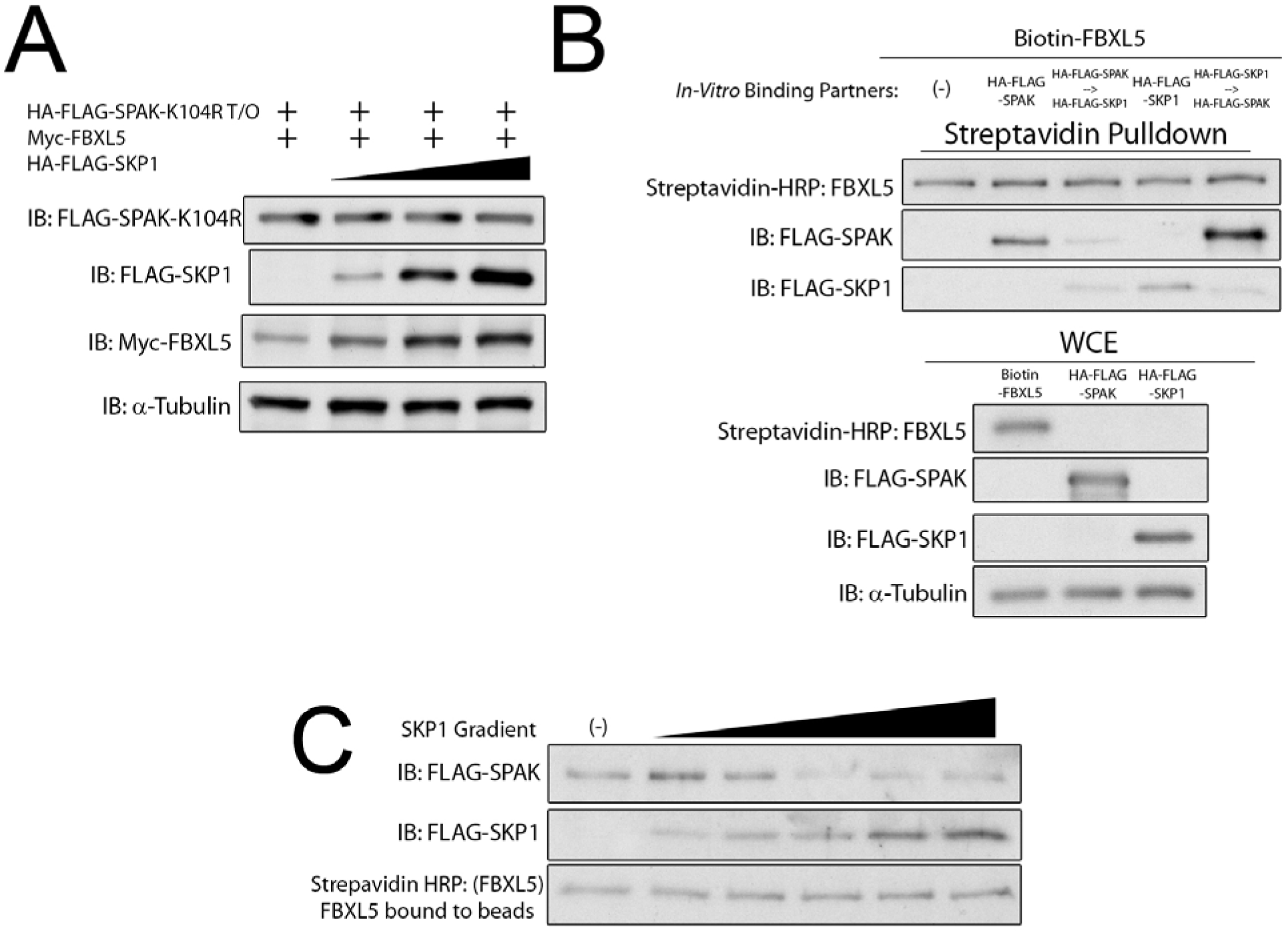
SPAK and SKP1 compete for binding to FBXL5. (**A**) HEK293 cells stably expressing 3HA-3FLAG-tagged SPAK-K104R were co-transfected with 6Myc-tagged FBXL5 and increasing concentrations of 3HA-3FLAG-tagged SKP1. WCEs were blotted with antibodies against FLAG, c-Myc and tubulin-HRP. (**B**) An *in vitro* binding assay was performed using BioEase-FBXL5 bound to streptavidin beads and incubated with either 3HA-3FLAG-SPAK or 3HA-3FLAG-SKP1. Beads were then washed and incubated a second time with the other 3HA-3FLAG-tagged protein. Streptavidin pulldowns and WCEs were blotted with streptavidin-HRP, FLAG and tubulin-HRP. (**C**) An *in vitro* binding assay was conducted using BioEase-FBXL5 bound to streptavidin beads and pre-bound to 3HA-3FLAG-tagged SPAK. The beads were then incubated with increasing concentrations of 3HA-3FLAG-SKP1. Streptavidin pulldowns were blotted with streptavidin-HRP and antibody against FLAG.

## DISCUSSION

FBXL5 is a substrate adaptor for the SKP1-CUL1-Fbox (SCF) family of E3 ubiquitin ligases and plays a central role in regulating iron homeostasis by degrading IRPs. As FBXL5 stability is directly coupled to intracellular iron levels, FBXL5 acts as a key homeostatic switch that degrades IRPs in iron replete conditions while being degraded itself when cellular iron levels are low and thereby allowing IRPs to accumulate and regulate their downstream RNA targets. Although this switch is essential for intracellular iron homeostasis, the pathways and components involved in regulating the stability of FBXL5 are still poorly characterized. Here, we report that FBXL5 physically interacts with SPAK, a kinase previously demonstrated to play a central role in the cellular response to osmotic stress (17). Moreover, this interaction is important for the proper regulation or iron homeostasis as either #1: the depletion of SPAK by siRNA or #2: the expression of constitutively active or kinase-dead mutants of SPAK influences the abundance, stability, and ubiquitination state of FBXL5 as well as the abundance of the iron storage protein ferritin, an important downstream effector of FBXL5. Similarly, an FBXL5 mutant whose binding to SPAK is impaired displays a significant reduction in stability. Together, these data support a novel role for SPAK in regulating iron homeostasis through its interaction with FBXL5.

Although these findings suggest that that SPAK–FBXL5 interaction can influence FBXL5 stability and iron homeostasis, the exact mechanism by which it does so remains unclear. Two experimental observations provide a potential framework for understanding this regulatory event. First, this interaction is iron-regulated and occurs preferentially in iron-replete conditions. Second, SPAK and SKP1 associate with FBXL5 in a competitive and mutually exclusive manner via an overlapping binding site on FBXL5. This suggests that SPAK associates with free FBXL5 and regulates the stability of a discrete pool of FBXL5 that is not in complex with the rest of the SCF complex under high iron conditions. At this point, both the function of this pool of FBXL5 as well as the substrates of SPAK that mediate this function have not been delineated. However, given that disruption of this interaction disturbs cellular iron homeostasis and that SPAK mutants that vary in kinase activity differentially affect iron homeostasis, these data are consistent with the SPAK–FBXL5 complex playing an important physiological role that will be important to elucidate in the future. Although we have focused on characterizing a potential role for SPAK in regulating iron homeostasis via its association with FBXL5, a second plausible possibility for the relevance of this interaction is that FBXL5 regulates SPAK activity and/or function in the context of the osmotic stress response. In the canonical SPAK signaling cascade, osmotic stress activates the WNK family of kinases which phosphorylate and activate SPAK and its close homologue OSR1 and which in turn regulate the activity of NKCC1, a sodium potassium chloride transporter that maintains electrolyte and fluid balance in cells (17). Future work examining whether changes in FBXL5 activity specifically or iron homeostasis more generally can influence SPAK signaling in these other physiological contexts will be of great interest moving forward. The identification of SPAK as a key regulator in both pathways may also suggest that crosstalk between iron homeostatic and osmotic signaling pathways is physiologically important.

## Acknowledgments

This work was supported in part by NIH grants GM089778 and GM112763 to J.A.W. D.N.P. was supported in part by the Ruth L. Kirschstein National Research Service Award GM007185 and the UCLA Dissertation Year Fellowship.

## Author Contributions

D.N.P. and J.A.W. conceived the experiments. D.N.P. and A.A.V. performed all experiments. A.A.V. provided proteomics analysis. C.Y. and M.S. assisted with experiments. D.N.P. and J.A.W. analyzed the data and wrote the manuscript. All authors edited the manuscript.

